# Phenotypic shifts of tumor associated macrophages and STAT3 mediated suppression of myeloid derived suppressor cells drive sensitization of HER2^+^ tumor immunity

**DOI:** 10.1101/2021.01.29.428708

**Authors:** Dimitrios N. Sidiropoulos, Christine Rafie, Brian J. Christmas, Emily F. Davis-Marcisak, Gaurav Sharma, Emma Bigelow, Anuj Gupta, Srinivasan Yegnasubramanian, Vered Stearns, Roisin M. Connolly, Daria A. Gaykalova, Luciane T. Kagohara, Elizabeth M. Jaffee, Elana J. Fertig, Evanthia T. Roussos Torres

## Abstract

Understanding how novel therapeutic combinations alter solid tumor microenvironments (TME) in immunosuppressive tumors such as breast cancer is essential to improve their responses to immune checkpoint inhibitors (ICIs). Entinostat, an oral histone deacetylase inhibitor (HDACi), has been shown to improve responses to ICIs in various tumor models with immunosuppressive TMEs, but the precise alterations induced by entinostat and mechanisms of synergy with ICIs remain unknown. Here, we employ single-cell RNA-sequencing on HER2 overexpressing breast tumors from mice treated with entinostat + ICIs to characterize these changes across cell types in the TME. This analysis demonstrates that treatment with entinostat induces a shift from a pro-tumor to an anti-tumor TME signature characterized predominantly by changes in the myeloid cells. Notably, myeloid-derived suppressor cells (MDSCs) are shifted toward the less suppressive granulocytic phenotype in association with reduced signaling through the STAT3 pathway. In addition, tumor-associated macrophages are shifted toward an anti-tumor M1 phenotype by epigenetic reprogramming. Overall, these entinostat-induced TME changes reduce immunosuppression and increase mechanisms of tumor cell killing to improve ICI responses and broaden the population of patients who could potentially benefit from immunotherapy.

## INTRODUCTION

Immune checkpoint inhibitors (ICIs) against programmed death receptor 1 (PD-1) and cytotoxic T-lymphocyte-associated protein 4 (CTLA-4), have revolutionized the treatment of immunogenic tumors that can mount a robust immune response due to the intrinsic presence of cytotoxic T cells. ICIs prevent inhibitory signaling that results in dampened cytotoxic T cell activity and permits a durable anti-tumor response in previously incurable cancers (Hodi et al. 2010; Wolchok et al. 2013). Nonetheless, tumors with a tumor microenvironment (TME) dominated by immunosuppressive cell types often do not respond to ICIs (Hodi et al., 2010). The HER2^+^ breast TME has been shown to include many cell types that prevent infiltration and activation of anti-tumor immune cells, including immunosuppressive cells such as T regulatory cells (Tregs), myeloid derived suppressor cells (MDSCs), and cancer associated fibroblasts (CAFs) (Umansky et al. 2013). These cells secrete suppressive cytokines and up-regulate the expression of immune checkpoint on T cells that bind to ligands in the TME and halt cytotoxic T cell activity (Paluskievicz et al. 2019). Although ICIs have been generally unsuccessful in these tumors, additional drugs have been explored in combination with ICIs to convert suppressive TMEs into immune permissive environments (Yong Li et al. 2018; Wall et al. 2020).

Epigenetic modulatory drugs, such as entinostat, are agents that have been shown to reprogram the TME to alter the abundance and immunosuppressive function of MDSCs to enhance the efficacy of ICIs in murine models of breast and pancreatic cancer (Christmas et al. 2018; Kim et al. 2014). However, the specific mechanisms of myeloid cell modulation by entinostat is not fully understood. Due to the heterogeneity of cell types altered by entinostat, single-cell RNA-sequencing (scRNAseq) is uniquely positioned to identify both the molecular and cellular changes induced by entinostat within MDSCs as well as all other cell types within tumors. We therefore employed scRNAseq to comprehensively characterize the signals altered in cell types within the TME following treatment with entinostat, and with entinostat + ICIs in the NeuN HER2-tolerized NT2.5 syngeneic model of breast cancer (Reilly et al. 2000; Mace et al. 1998). Our data demonstrate that entinostat regulates MDSCs in NeuN tumors through altered phosphorylation of STAT3 and regulation of IkBz. We also identified that M1 and M2 tumor associated macrophages (TAMs) are altered by combination treatment with entinostat and ICIs and contribute to transformation of the TME to support tumor cell killing. These findings point to a new mechanism by which entinostat sensitizes tumors to ICIs.

## RESULTS

### Single-cell RNA-sequencing reveals cellular composition within the breast tumor microenvironment of NeuN mice

To evaluate the molecular and cellular determinants of entinostat’s impact on the immunosuppressive TME, we analyzed tumors derived from NeuN mice. This tolerized HER2 overexpressing model of breast cancer mimics the non-immunogenic TME observed in breast cancer patients, making it an ideal model to study poor responders to currently available ICIs. We performed scRNAseq on whole tumors following treatment with entinostat alone or in combination with ICIs to characterize the pathways impacted by entinostat and their further interaction with immunotherapy treatment. Tumor were implanted and mice were treated for 3 weeks with vehicle control (V), entinostat alone (E) or with entinostat in combination with anti-PD-1 (EP), with anti-CTLA-4 (EC), or the triple combination (EPC) (Figure 1A). After 3 weeks, tumors were harvested, processed into single cell suspension, and scRNAseq was performed using the 10x Genomics platform. Individual cells from each tumor were annotated into cell types by clustering the RNA expression profiles then annotated and assessing the expression of known cell type markers (Supp. Table 1). This analysis grossly characterized cells into 5 cell populations that included cancer cells, CAFs, lymphoid cells, monocytes/macrophages, and MDSCs. Each of these cell types were represented in tumors from all treatment conditions (Figures 1B, 1C, Supp. Table 1).

Cellular abundances among the five broad cell types identified in the NeuN model did not vary significantly between treatment groups (Figure 1D, 1E). Still, these data enabled further investigation of cell type specific transcriptional differences and cell state transitions associated with each treatment. Notably, we found significantly increased expression of *Il2,* and lymphocyte maturation factor *Ly6c2* in cancer cells from mice treated with entinostat vs. vehicle (Supp. Figure 1A). We performed further gene set analysis to determine which pathways were enriched with the gene expression changes between treatments in each cell type. This analysis identified enrichment of KEGG pathways related to immune response, including statistically significant overexpression of the genes involved in antigen processing and presentation in the cancer cells *(q=0.004)* (Supp. Figure 1B) and chemokine signaling in the CAFs *(q=0.048)* (Supp. Figure 1C). These results suggest that treatment with entinostat may prime NeuN tumors and improve response to combinatorial treatment of entinostat combined with ICIs. Moreover, T regulatory cells dominate the TME of the lymphocyte cluster, which is consistent with the immunosuppressive nature of the NeuN model (Supp. Figures 2A, 2B, Supp. Table 1). Treatment with entinostat + ICIs trended toward increased Tregs (Supp. Figure 2C). The myeloid cells (MDSCs and macrophages) were consistently the most abundant cell type in our scRNAseq data from the NeuN model (Figure 1E), further supporting the significant role these cell types play in regulating immune response in breast cancer. Therefore, we focused our in-depth analysis of the molecular changes induced by entinostat and ICI treatment on MDSCs and macrophages.

**Figure 1.**
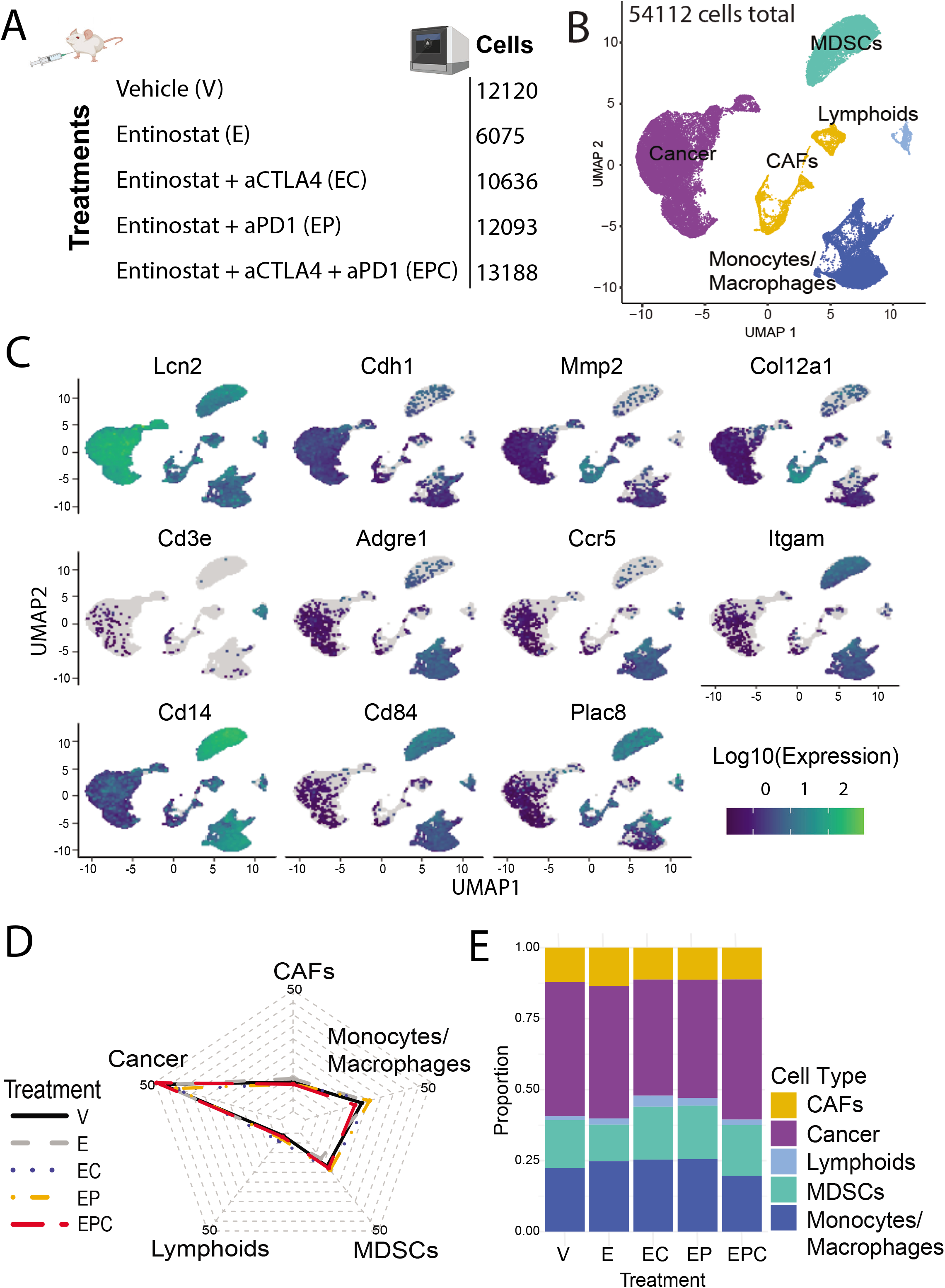
Single cell transcriptional atlas of NeuN tumors treated with HDACi alone or in combination with ICI. **A,** Treatment groups. **B,** UMAP with each cell partition annotated by its corresponding cell type determined from top differentially expressed markers. **C**, Representative gene markers. **D**, Radar plot and stacked bar plot (**E**) of broad cell type composition normalized to total cell count from each group.

### Myeloid cells predominate the breast immune TME

We performed deeper clustering analysis of the MDSC and monocyte/macrophage clusters to further characterize these cell populations (Figure 2A. 2B, Supp. Table 1). Expression of functional genes such as *Tnf, Itgax, Flt3, Itgae, Ccr7, Cx3cr1, Ccr2, Ccr5, C1qc,* and *Arg1* was used define sub-populations of cells within the identified MDSC and macrophage clusters (Figure 2A). Our analysis revealed 3656 monocytic-MDSCs (M-MDSCs), 5803 granulocytic-MDSCs (G-MDSCs), 307 dendritic cells (DC), 1167 monocytes, and 11093 macrophages (TAMs) (Supp. Table 1). Since macrophages can be either pro-tumor or anti-tumor in function, we further delineated the TAMs cluster using gene expression signatures representative of M1 (anti-tumor) and M2 (pro-tumor) TAMs (Figure 2C). When visualizing the distribution of these myeloid subsets among the various treatments, treatment with EPC induced a shift from a TME dominated by M-MDSCs and M2 macrophages to one primarily composed of G-MDSCs and M1 macrophages (Figure 2D).

**Figure 2.**
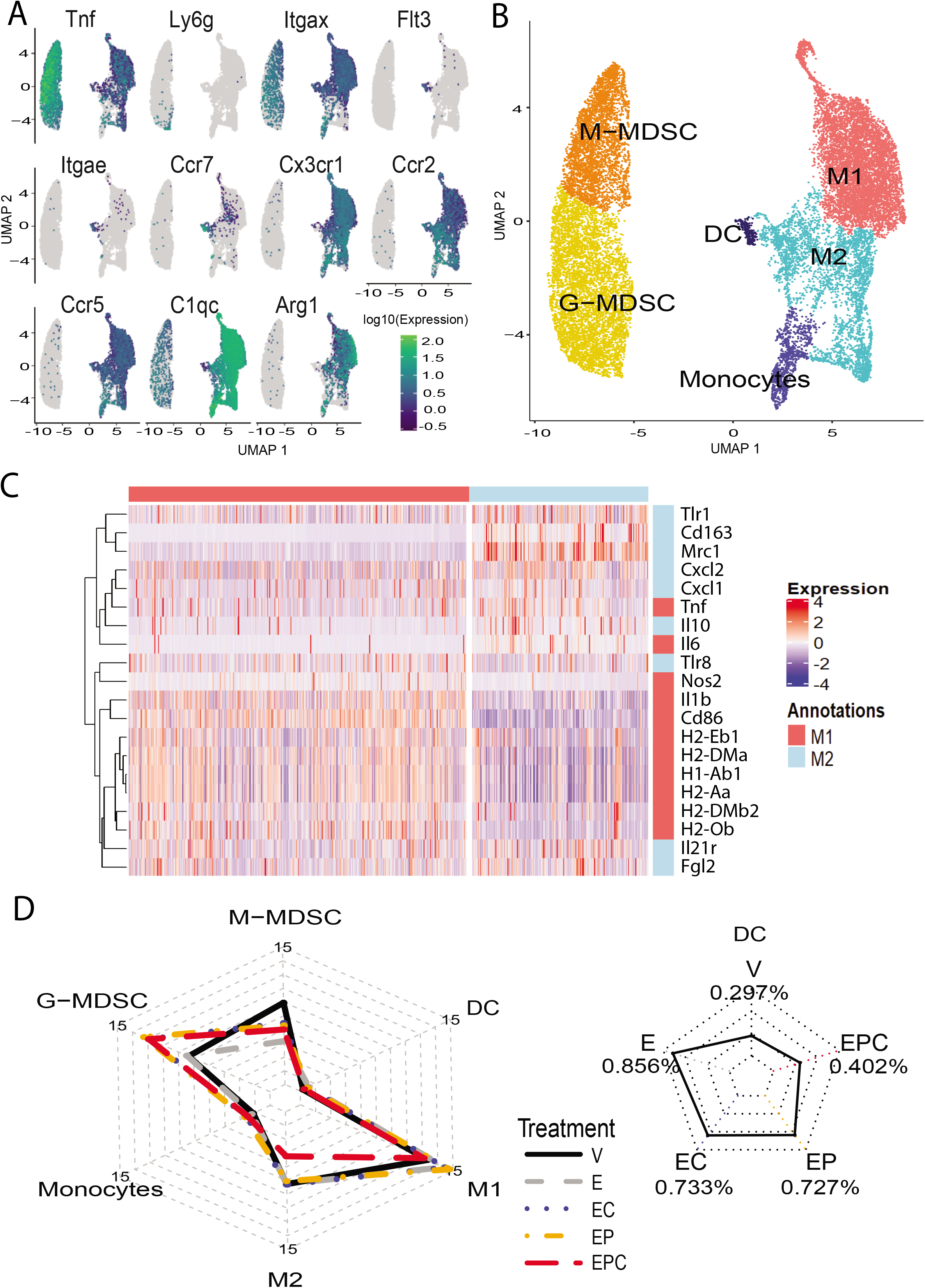
Pooling myeloid-derived populations reveals distinct subpopulations that are differentially abundant across treatments. **A,** UMAPs of representative myeloid cell subtype markers. **B**, Annotated clusters projected onto the UMAP of the myeloid subset population. **C,** Pearson-clustered heatmap of TAM cells and curated canonical markers depicting matching expressions of M1 and M2 clusters in the scRNAseq data. **D,** Radar plot of myeloid cell type composition within the TME normalized to total cell count from each treatment group (left) and radar plot of DC composition by treatment (right).

Ratios of G/M-MDSCs and M1/M2 macrophages reflect suppressive function and pro vs. anti-tumor response, respectively (Je-In Youn et al. 2012; Macciò et al. 2020). We found an elevated M1/M2 macrophage ratio with combination treatment and elevated G/M-MDSC ratio with entinostat treatment (Figures 3A, 3B), which was corroborated by changes in flow cytometry analysis of tumors from NeuN treated mice (Figures 3C, 3D). Gene expression and flow data suggest an entinostat-mediated enrichment of the anti-tumor myeloid cells. An increased G/M-MDSC ratio with entinostat treatment suggests a shift to a more immunogenic TME, as G-MDSCs are less suppressive than M-MDSCs (Veglia et al. 2018; Kumar et al. 2017). In addition, an increased M1/M2 macrophage ratio suggests a more robust anti-tumor response following combination treatment. Both of these shifts in cell phenotypes support a sensitized TME with treatment of entinostat and ICIs.

**Figure 3.**
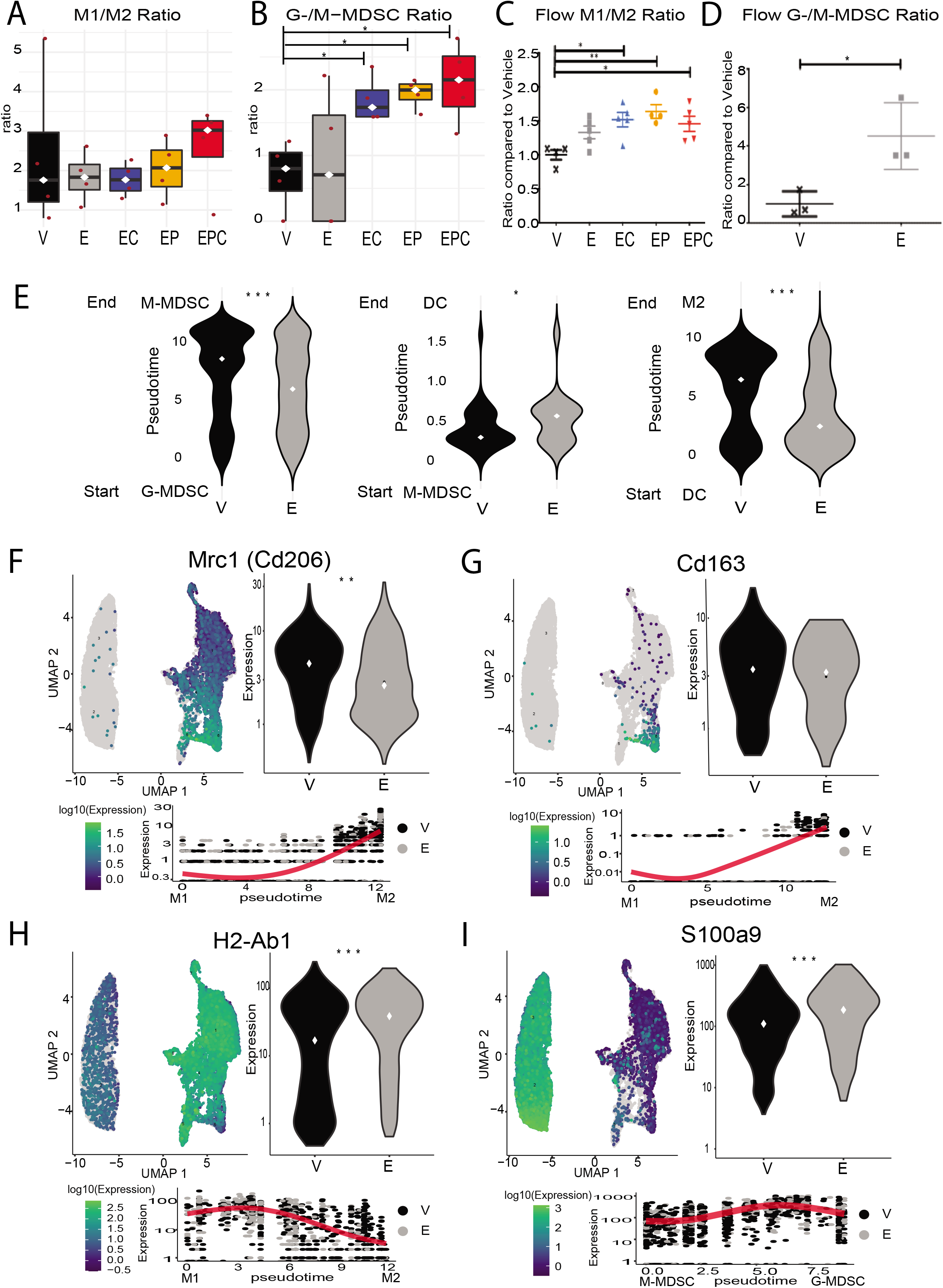
Treatment with entinostat and ICIs induces anti-tumor shifts in myeloid populations in the breast immune TME. **A-B,** M1:M2 ratio **(A)** and G-MDSC:M-MDSC ratio (**B)** by tumor graphed according to the single cell RNAseq data, each red dot corresponding to one tumor, significance calculated using Welch Two Sample t-test. **C-D**, Flow cytometry analysis of G:M-MDSC ratios (**C**) and M1:M2 macrophage ratios (**D**) of breast tumors from entinostat and ICI treated NeuN mice. Each dot represents one mouse and each bar represents mean +/− SEM; n=3–5 mice/group. Significance was determined by a one-way ANOVA with Tukey multiple comparisons test. **E,** Violin plots comparing Vehicle and Entinostat pseudotime weights for M-MDSCs (Left), DCs (Middle), and M2 macrophages (Right), using G-MDSCs as the starting node reference point. **F-I** UMAPs of myeloid cells colored by expressions of representative differentially expressed genes *Mrc1, Cd163, H2-Ab1,* and *S100a9* accompanied by violin plots of their gene expression comparing Vehicle and Entinostat and scatter plots showing expression of each gene by pseudotime. Gene expression statistics determined using monocle3 negative binomial model. Pseudotime statistics were calculated using a Wilcoxon rank sum exact test. Statistically significant P values are shown as follows: * P < 0.05; ** P < 0.01; *** P < 0.001.

### Molecular changes in myeloid cell subsets promote less suppressive G-MDSCs and antitumor M1 macrophages by entinostat and ICIs

We performed pseudotime analysis to evaluate if treatments impact cell state transitions associated with changes in immunosuppression from myeloid and macrophage populations. When measuring differential weights of pseudotime analysis, we observed that there are higher weights and thus higher representation of M-MDSCs and M2 macrophages in vehicle relative to entinostat treated tumors (Figure 3E) which is representative of a state change reducing immunosuppressive cellular function. Further evaluation of the pseudotime analysis of cells in the TAM cell cluster reveals a branch point that suggests cells split off into two cell clusters from the DCs: an M1 and an M2 TAM sub-cluster.

We evaluated the expression of some of the significantly differentially expressed genes (e.g., *Mrc1, Cd163, H2-Ab1* and *S100a9,* Supplemental File 1) that are known to play a role in immunosuppressive function and phenotype switching. We correlated the molecular changes with pseudotime trajectories to determine if changes in gene expression was correlated with a cell phenotype switch. For example, TAM M2 markers *Mrc1/CD206* and *Cd163* are significantly decreased upon entinostat treatment (Figure 3F, *q=0.011,* 3G, *q value* over FDR threshold, *p=4.36e-06).* This decrease correlates with an M2 to M1 phenotype switch, suggesting that these genes are driving the shift in the TAM trajectory. Expression of *H2-Ab1,* which encodes MHC-II, is significantly upregulated with entinostat treatment in M1 macrophages and correlates with the pseudotime trajectory predicting a shift from M2 to M1 TAMs (Figure 3H, *q=7.72e-08).* Similarly, significantly altered gene expression in MDSCs from entinostat treated tumors revealed an upregulation of *S100a9,* which is known to be more highly expressed in G-MDSCs (Veglia et al. 2018) (Figure 3I, *q=0.000266).* This suggests that upregulation of *S100a9* induced by entinostat represents a shift from an M-MDSC to a G-MDSC phenotype and is potentially driving the increased ratio of G/M-MDSCs. Taken together, these results suggest that M2 macrophages and G-MDSCs are most significantly affected by treatment with entinostat and may be driving the transition to an immuno-permissive TME.

### Entinostat treatment dampens suppressive capabilities of MDSCs by altering STAT3 signaling

The scRNAseq data suggests that entinostat induces cell state transitions from G-MDSCs into less immunosuppressive states to promote ICI sensitivity. Our previous functional data shows entinostat specifically decreases the suppressive function of G-MDSCs and suggests that this is due to the altered phosphorylation of STAT3 in G-MDSCs (Christmas et al. 2018). To investigate whether this mechanism is associated with the cell state transitions observed in the single cell analysis, we first validated functional changes induced by entinostat in an MDSC-like cell line (J774M cells). Consistent with findings from intratumoral MDSCs, we observed a significantly decreased capacity to suppress T cell proliferation upon entinostat treatment (Suppl. Figure 4A-B), and a decreased IL-6 induced STAT3 phosphorylation (Suppl. Figure 4C-D). We then performed chromatin immunoprecipitation sequencing (ChIP-seq) on an MDSC-like cell line (J774M cells) treated with entinostat using a pSTAT3-specific antibody. We analyzed the ChIP-seq data for peaks that were unique to untreated vs. entinostat treated MDSCs to reveal differential pSTAT3 binding induced by entinostat. We hypothesized that pSTAT3 binding observed only in entinostat-treated MDSCs would be associated with the expression of genes responsible for MDSC suppression changes. We identified pSTAT3 binding peaks within the promoters of 35 unique genes in entinostat treated compared to untreated MDSCs, which exhibited pSTAT3 binding peaks within the promoters of only 6 unique genes (Suppl. Table 2). We performed an integrated analysis of the ChIP-seq and scRNAseq data to determine if the 35 genes whose promoters were bound by pSTAT3 in entinostat-treated MDSCs were differentially expressed in G vs M-MDSCs. We found that gene expression of pSTAT3 target genes were enriched in G-MDSCs relative to M-MDSCs *(p=0.0245,* Figure 4A), suggesting that the targets in the *in vitro* model are consistent with pSTAT3 activation in G-MDSC cell populations *in vivo.*

**Figure 4.**
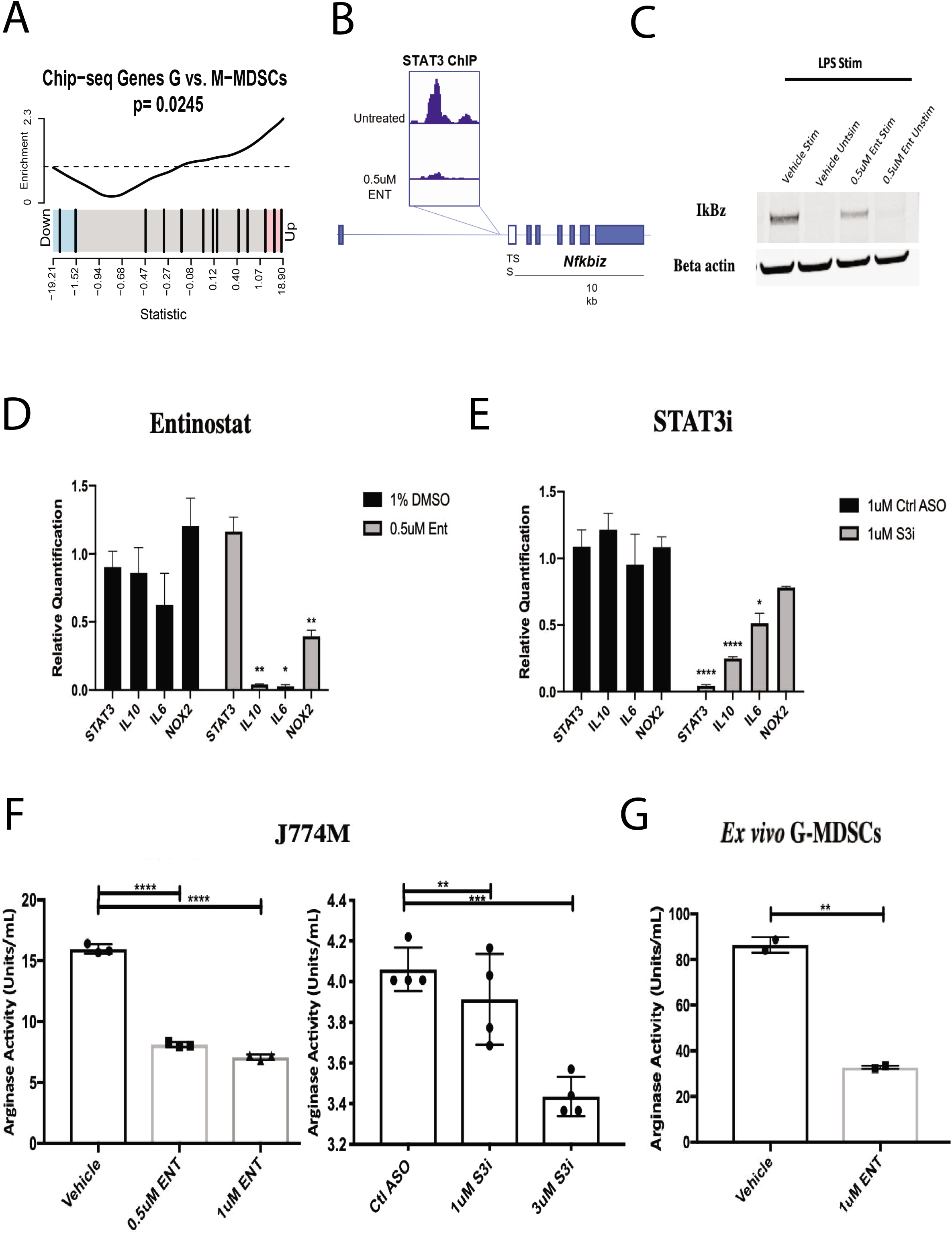
Effects of entinostat are mediated by STAT3 modulation. **A,** Barcode plot that ranks the ENT-altered STAT3 target genes and shows they are more enriched in G-MDSCs over M-MDSCs. **B,** ChIP-seq was performed on J774M cells treated for 16 hours *in vitro* with entinostat, using a phosphorylated STAT3 antibody for pulldown. ENT reduced pSTAT3 binding on downstream promoters including *Nfkbiz* (see table in supplemental for all peaks). **C**, J774M cells pre-treated with entinostat for 16 hours were stimulated with LPS and lysed. Western blot depicts lysates probed for IkBz and beta-actin. **D and E**, qPCR of inflammatory genes implicated by ChIP-seq was performed on J774M cells treated with entinostat for 16 hours or STAT3 ASO for 72 hours (n=3 biological replicates/group). **F and G**, Arginase enzyme activity in J774M cells was measured after 16 hours of treatment with ENT or 72 hours STAT3 ASO. Each dot represents one biological replicate and each bar represents mean ± SEM. **H**, Ly6G^+^ G-MDSCs were isolated from tumors of untreated neuN mice and treated with ENT ex vivo for 16 hours in tumor conditioned media before arginase activity was measured. Each dot represents G-MDSCs pooled from two tumors and each bar represents mean ± SEM. Statistically significant p values are aollows: *p<0.05, **p<0.01, ***p<0.001, ****p<0.0001.

Moreover, entinostat treatment prevented the binding of pSTAT3 to the promoter of *Nfkbiz* (Figure 4B), a known regulator of pro- and anti-inflammatory cytokines in myeloid cells (Horber et al. 2016). These results suggest that entinostat treatment restricts a STAT3-mediated inflammatory response and induces an alternate gene profile in these cells. To confirm that this decrease in pSTAT3 binding to the *Nfkbiz* promoter resulted in a subsequent decrease in its downstream protein, we performed Western blots probing for IkBz after entinostat treatment and found a decrease in IkBz production (Figure 4C). These results indicate that entinostat treatment engages alternative inflammatory STAT3 signaling in MDSCs.

To determine if changes in IkBz led to decreased production of genes responsible for suppressive function, we looked at expression of immunosuppressive cytokines following treatment with either entinostat or a STAT3 antisense oligonucleotide inhibitor (STAT3i ASO, AstraZeneca) which is known to specifically decrease the phosphorylation of STAT3. We found that treatment with entinostat led to decreased production of *IL-10, IL-6,* and *NOX2* while treatment with STAT3i resulted in decreased production of *IL-10* and *IL-6* (Figure 4D, 4E). At the protein level, we found that entinostat and STAT3i ASO decreased the activity of Arg-1 protein in J774M cells (Figure 4F). Similarly, entinostat decreased Arg-1 activity in intratumoral G-MDSCs (Figure 4G). Altogether, these data show that entinostat promotes a less suppressive MDSC phenotype via its effect on phosphorylation of STAT3 in MDSCs.

In summary, this data suggests that entinostat treatment promotes tumor killing mechanisms via shifts from pro-tumor myeloid cells towards professional antigen-presenting cells (i.e. DCs and M1 TAMs) and reduces immunosuppression via shifts in MDSC cell subtypes and function (Figure 5). TAMs are skewed toward an M1 anti-tumor phenotype via decreased expression of *Mrc1* and *Cd163* and increased *H2-Ab1.* Myeloid DCs are enriched as evidenced by the expression of *Flt3, Itgae, Ccr7, Btla,* and *MHCI/II.* We hypothesize that this ultimately contributes to T cell recruitment, infiltration and improved cytotoxicity (Figure 5). In addition, entinostat targets MDSCs and skews them toward a less suppressive G-MDSC phenotype, compounded by decreased pSTAT3 by entinostat, which leads to decreased production of suppressive cytokines. These changes all drive the non-immunogenic breast tumor to become more immunogenic and thus more likely to respond to ICIs to promote tumor killing.

**Figure 5.**
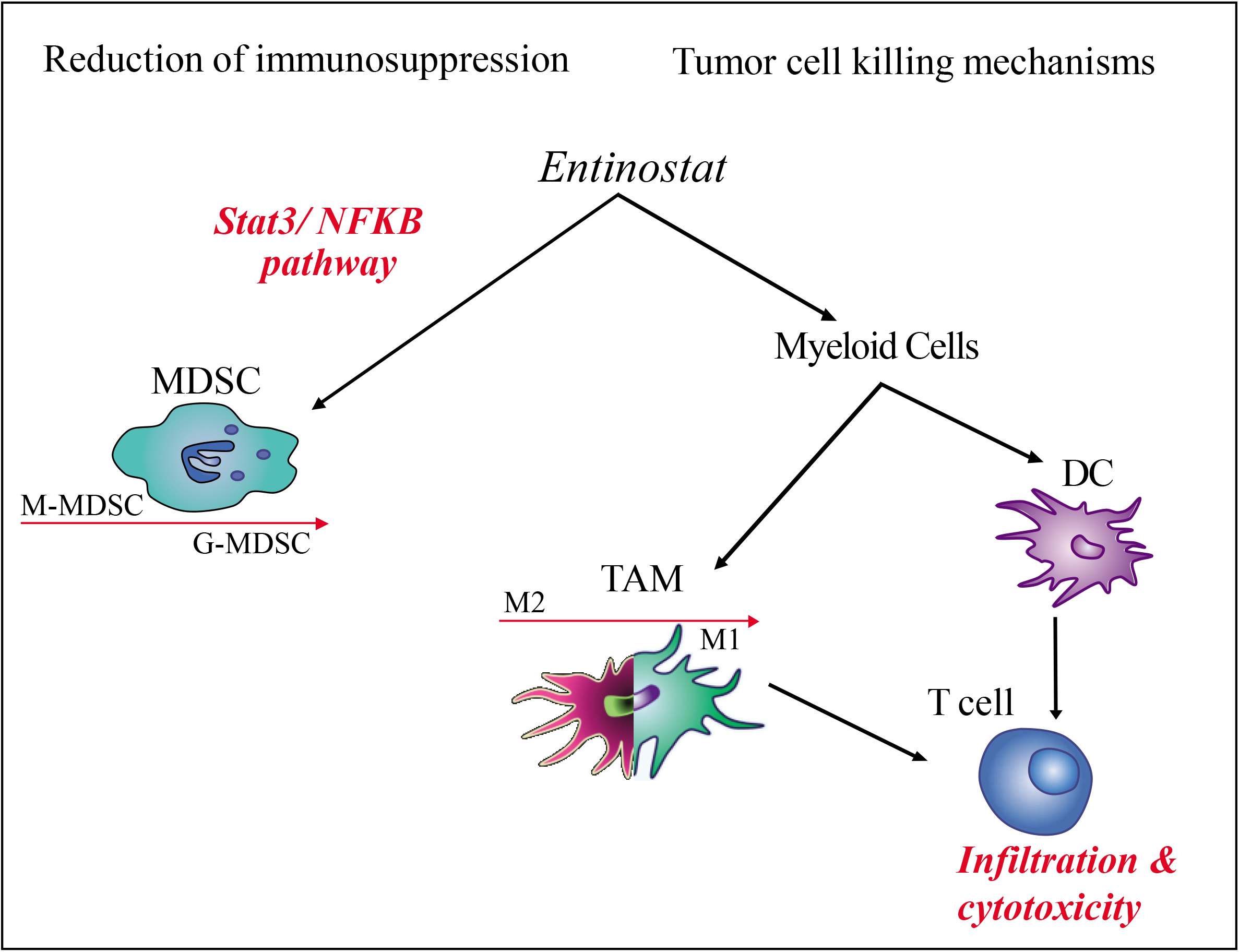
Proposed model of molecular mechanisms affected by treatment in the myeloid lineage of the TME. Based on our results, Entinostat promotes an anti-tumor shift in the TME. This is likely accomplished by polarizing MDSCs through altered STAT3 signaling, expanding myeloid DCs, and in combination with ICIs polarizing TAMs from an M2 towards an M1-like phenotype.

## DISCUSSION

This study provides an in-depth characterization of the cellular composition and transcriptional changes in the TME from the NeuN murine model of breast cancer treated with entinostat in combination with ICIs. Using scRNAseq, we delineated 5 different cell populations within the TME and investigated differential gene expression in the two most abundant immune cell populations, MDSCs and TAMs, as well as differences in cellular subtypes in response to treatment with entinostat and ICIs. Treatment with entinostat combined with ICIs skewed cell populations of myeloid lineage to favor G-MDSCs and M1 macrophages. The less suppressive nature of G-MDSCs and the pro-inflammatory characteristics of M1 macrophages suggests treatment with entinostat + ICIs promotes an anti-tumor TME. We also uncovered specific gene targets, such as *Mrc1, Cd163,* and *H2-Ab1,* that could contribute toward the skewing of TAMs to M1 phenotypes. Additionally, we found changes in *S100a9* and *Nfkbiz* expression could contribute to decreased MDSC suppression. These findings reveal immune cell subsets and novel targets within the TME that, upon further validation, will inform therapeutic combinations with ICIs to help improve response rates for patients with non-immunogenic tumors.

One of the novelties of this work is our innovative use of scRNAseq to interrogate tumors from different therapeutic treatment groups. Analysis of these data helped us identify cell types such as G/M-MDSCs and M1/M2 macrophages, as well as pathways within tumor cells and CAFs that are all targeted by the combination of entinostat + ICIs and alter the TME into one that is more likely to respond to ICIs. The combination of entinostat + ICIs induced the most significant changes in MDSCs and cells of monocyte lineage, such as DCs and TAMs. MDSCs accumulate in the TME and interfere with anti-tumor immune responses (Ostrand-Rosenberg et al. 2018). Thus, it is not surprising that our analysis revealed approximately a 1:1 ratio of myeloid cells to cancer cells in all treatment groups, further emphasizing the importance of the myeloid immune cell populations in breast tumors. Differential expression analysis between treatment groups allowed us to pinpoint molecular mechanisms underlying dynamic shifts between MDSC subtypes in the TME.

Between the two studied MDSC subtypes, G-MDSC and M-MDSC, the latter is the more suppressive type when assessed on a per-cell basis and their mechanisms of immunosuppression more prolonged (Veglia et al. 2018; Kumar et al. 2017). Thus, various efforts have been made to pharmacologically target them in order to increase ICI efficacy. For example, blockade of STAT3 via a TLR9 agonist, has been shown to inhibit MDSC suppression (Qifang Zhang et al. 2016). Our integrated analysis of data from scRNAseq with ChIP-seq supports our previous findings that entinostat treatment induces changes within the STAT3 signaling pathway (Christmas et al. 2018) and demonstrates further that this leads to alterations in MDSC function by decreasing immunosuppressive cytokine production. We show one mechanism by which entinostat alters MDSC suppressive function is through decreased phosphorylation of STAT3. Decreased pSTAT3 then leads to decreased transcription of *Nfkbiz,* and subsequently decreases the expression of IkBz (a downstream effector of the NFKB pathway) as well as the immunosuppressive cytokines IL6 and IL10. This is consistent with a previous report showing that knock out of *Nfkbiz* in macrophages results in a decrease in IL6 and IL10 production and promotes a more M1 phenotype (Horber et al. 2016). Additionally, others have shown that: (1) STAT3 binds the promoter of *Nfkbiz* in the same location identified by our ChIP-seq analysis (Muromoto et al. 2019) and (2) knockout of STAT3 results in a decrease in IkBz expression (Okuma et al. 2013). Taken together, these data strongly suggest that STAT3 signaling plays a role in regulating IkBz expression and its subsequent regulation of the inflammatory state of macrophages.

The single cell data in this study also elucidate the impact of treatment on myeloid and macrophage phenotype switching within the breast TME. Myeloid cells have been described as a fluid cell type recruited by tumor cells, and then reprogrammed to inhibit the anti-tumor response (Gabrilovich et al. 2012). Specifically, intratumoral MDSCs can be shifted from the G-to M-MDSC phenotype and differentiate into TAMs and DCs and possibly support an anti-tumor response (Gabrilovich et al. 2012). We previously reported an increase in tumor infiltrating G-MDSCs without a notable shift in M-MDSCs (Christmas et al. 2018). Our current study suggests treatment with entinostat could promote the phenotypic switch and subsequent differentiation into TAMs and DCs. In untreated tumors, we observed accumulation of M-MDSCs and M2-TAMs that harbor a pro-tumorigenic signature. This was confirmed using differential abundance and pseudotime weights showing that untreated tumors have significantly more myeloid cells progressing along M-MDSC and M2-TAM differentiation axes, whereas myeloid cells from entinostat treated tumors are shifted toward G-MDSCs, DCs and M1-TAMs. An additional example of cell phenotype switching in our analysis is the suggested polarization of tumor-associated macrophages (TAMs) by entinostat. We observed enrichment of M1 TAMs in tumors treated with combination therapy indicative that epigenetic modulation alone is not sufficient to polarize M2 TAMs to M1, and the addition of ICIs to entinostat seems to support M1 TAM polarization. Treatment with entinostat alone downregulates canonical markers of M2 TAMs—*Mrc1/CD206* and *Cd163,* supportive of the hypothesis that entinostat initiates induction of TAM polarization toward an M1 phenotype.

Altogether, our analysis across cell types leads us to propose a model that entinostat has the two-fold function of reduction of immunosuppression and tumor cell killing (Figure 5). Notably, the inferred myeloid cell state transitions from our scRNA-seq data lead to a central hypothesis that terminal differentiation into DCs and TAMs and further switching from an M2 to M1 phenotype improves the anti-tumor response in breast cancer, further supported by similar findings in sarcoma (Devalaraja et al. 2020). In this model, immunosuppression is decreased following altered binding of pSTAT3 to promotors controlling downstream regulators of the NFkB pathway and effectors of suppressive signaling such as IL-6 and IL-10. These data suggest that pSTAT3 mediated skewing of the MDSC phenotype from M-MDSC to G-MDSCs is an area of future mechanistic study and evaluation in human tumors treated with entinostat, nivolumab, and ipilimumab in a clinical trial (NCT02453620) mirroring the preclinical treatments presented in this study. While entinostat may promote these cell state transitions directly, the changes it induces in in chemokine signaling and antigen processing and presentation pathways in tumor cells may further introduce inter-cellular interactions that promote the observed cell state transitions in the myeloid compartment. Ultimately, our findings also provide rationale for investigation of more specific cellular targets within the breast TME such as M2 macrophage or G-MDSCs as a foundation to evaluate new treatment modalities to sensitize breast cancers to immunotherapy.

## METHODS

### Mice and cell lines

Animals were kept in pathogen-free conditions and were treated in accordance with institutional and American Association of Laboratory Animal Committee policies. NeuN mice were originally from W. Muller McMaster University, Hamilton, Ontario, Canada and overexpress HER2 via the mouse mammary tumor virus (MMTV) promotor. Colonies were renewed yearly from Jackson labs and bred in house by brother/sister mating. NT2.5 cells were derived from spontaneous HER2 overexpressing mammary tumors growing in female NeuN mice. *In vitro* cell lines were established and authenticated as previously described (Chunwan Lu et al., 2016). Culture conditions for NT2.5 cells were as follows: 37°C, 5% CO2 in RPMI 1640 (Gibco, cat. 11875-093) supplemented with 20% fetal bovine serum (Gemini, cat. 100-106), 1.2% HEPES buffer (Gibco, cat. 15630-080), 1% L-glutamine (Gibco, cat. 25030-081), 1% MEM non-essential amino acids (Gibco, cat. 11140-050), 0.5% penicillin streptomycin (Gibco, cat. 15140-122), 1% sodium pyruvate (Sigma, cat. S8636), 0.2% insulin (NovoLog, cat. U-100), 0.02% gentamicin (Sigma, cat. G1397). J774M cells have been previously established as a MDSC cell line (Lu et al 2016) and were kindly provided for use in this investigation by Dr. Kebin Liu. J774M cell line was developed by FACS sorting CD11b^+^ Gr1^+^ cells from the ATCC macrophage line J774A.1 and have previously been shown to exhibit immunosuppressive functions like MDSCs (Lu et al. 2016; Christmas et al. 2018). Culture conditions for J774M cells were as follows: 37°C, 5% CO2 in RPMI 1640 (Gibco, cat. 11875-093) supplemented with 10% fetal bovine serum (Gemini, cat. 100-106), 1.5% HEPES buffer (Gibco, cat. 15630-080), 1% L-glutamine (Gibco, cat. 25030-081), 1% MEM non-essential amino acids (Gibco, cat. 11140-050), 1% penicillin streptomycin (Gibco, cat. 15140-122), 1% sodium pyruvate (Sigma, cat. S8636), 0.0004% beta-mercaptoethanol (Sigma, cat. M3148). All cell lines were regularly tested for Mycoplasma every three months in accordance with laboratory policy.

### Tumor dissociation

To obtain single-cell suspensions from breast tumors, tumors were harvested, weighed, diced, and then dissociated using a tumor dissociation kit (Miltenyi Biotec, cat. 130-096-730) and the OctoDissociator (Miltenyi Biotec) per the manufacturer’s instructions. The 37C_m_TDK_2 program was used to dissociate tumors per the manufacturer’s instructions. Samples were filtered using a 40 mm cell strainer and red blood cells were lysed using ACK lysis buffer (Quality Biological, cat. 118-156-721). The resulting single-cell suspensions were used for subsequent isolation of specific immune cell types or flow cytometry as described below, or for scRNAseq following an additional step of dead cell removal using the MACS Dead Cell Removal Kit (Miltenyi Biotec).

### Immune cell isolation

G-MDSCs were isolated from single-cell suspensions from tumors following tumor dissociation using Miltenyi Biotec’s Myeloid-Derived Suppressor Cell Isolation Kit (cat. 130-094-538) according to the manufacturer’s protocol. Ly6G^+^ cells were positively selected to isolate G-MDSCs from Ly6G^-^ M-MDSCs and were passed through LS columns (Miltenyi Biotec, cat. 130-042-401) twice to increase purity. Eluted G-MDSCs were then used for downstream assays described below. CD8^+^ T cells were negatively isolated from spleens of c100+ Balb/c mice (above) by mashing spleens through 100 mm cell strainers, lysing red blood cells using ACK lysis buffer (Quality Biological, cat. 118-156-721), and isolating cells via the EasySep Mouse CD8^+^ Isolation Kit (StemCell, cat. 19853) per manufacturer’s instructions. CD8^+^ T cells were then used for the *in vitro* suppression assays described below.

### Flow cytometry

Isolated single-cell suspensions were washed and then incubated for 30 minutes with Live/Dead Near-IR (ThermoFisher, cat. L10119) according to the manufacturer’s protocol, followed by a 30-minute incubation with the appropriate flow cytometry antibodies. Samples were run on a CytoFLEX (Beckman Coulter) cytometer and analyzed using FlowJo (FlowJo LLC).

### Ex vivo G-MDSC culturing

For *ex vivo* experiments using intratumoral G-MDSCs, G-MDSCs were plated in tumor conditioned media after being isolated as described above. Tumor conditioned media is derived by culturing NT2.5 for 48 hours in the NT2.5 media described above and subsequent filtering to remove tumor cells.

### Arginase assay

Arginase activity was measured calorimetrically using Abcam’s Arginase Activity Assay Kit (cat. ab180877). For *ex vivo* studies, G-MDSCs were isolated from tumors, plated in tumor conditioned media with ENT or DMSO vehicle for 16 hours, and subsequently harvested. For *in vitro* studies, J774M were cultured with ENT or DMSO vehicle for 16 hours and subsequently harvested. In short, cells were lysed with the kit’s lysis buffer at 1×10^6^/1mL, and plated (500k cells per well) in duplicate in a flat-bottom, low-retention plate carefully to avoid bubble formation. Target samples were incubated for 20 minutes at 37°C with H2O2 substrate solution, while background wells were incubated with additional buffer. Standards were prepared per kit instructions, and the enzymatic reaction mixture was prepared and added to all wells. Raw absorbance values were immediately obtained over a 30-minute period using a plate reader (Molecular Devices SpectraMax M3) at OD=570nm at 37°C. Arginase Activity Units were then calculated from raw absorbance values. ΔOD (ΔOD= (OD2-ODbg2)-(OD1-ODbg1)) was used to obtain the nmol of H2O2 generated by arginase, collected from a standard curve of known H2O2 concentrations. Arginase activity is calculated as (B/ΔT*V)*D in units/mL, where B is amount of H2O2 from standard curve (nmol), V is the sample volume added into reaction well (mL), D is sample dilution factor. One unit of Arginase activity refers to the amount of arginase that will generate 1.0 nmol of H2O2 per minute at pH 8 at 37°C.

### In vitro suppression assay

J774M cells treated with ENT or DMSO vehicle for 16 hours were cocultured with stimulated T cells. T cell proliferation was measured via CFSE dilution. T cells were isolated from spleens of Balb/c mice as described above and subsequently labeled with CFSE (ThermoFisher, cat. C34554) per the manufacturer’s instructions. 2.5 x10^5^ CFSE-labeled CD8^+^ T cells were cultured with J774M cells at varying ratios (1:1 – 1:8 J774M:T cells) and anti-CD3/CD28 beads (ThermoFisher, cat. 11453D) at a bead-to-T cell ratio of 1:1 per the manufacturer’s instructions for T cell activation. T cells were allowed to proliferate for 52 hours. Subsequently, the cultures were harvested, stained with Live/Dead NIR (ThermoFisher, cat. L10119) as well as CD8 and analyzed via flow cytometric analysis as described above. Dilutions of initial CFSE were indications of T cell divisions, where fewer divisions indicated greater suppressive activity. All antibodies used are listed in Supplementary File 2.

### Western blots

For analysis of *in vivo* samples, G-MDSCs were isolated from the whole tumor of treated animals using Miltenyi Biotec’s MDSC Isolation Kit and pooled by treatment group. For *ex vivo* samples, G-MDSCs were isolated from untreated animals, cultured with ENT or DMSO vehicle in tumor conditioned media for 16 hours as described above, and subsequently harvested. For *in vitro* analysis, J774M cells were cultured with ENT or DMSO vehicle for 16 hours and subsequently harvested. Samples being used for phospho-STAT3 analysis were subsequently stimulated with IL-6 at 20 ng/μl for 25 minutes. Samples were lysed in RIPA buffer with added 1μM DTT, 1μM PMSF, and 1:100 protease/phosphatase inhibitor cocktail (Cell Signaling #5872S) and quantified by BCA (Pierce, #23225). Protein amounts were normalized between samples and run on a 4-12% Bis-Tris gels under denaturing conditions. The LiCor Odyssey developing and imaging system was used, and the following primary antibodies were diluted in Odyssey Blocking Buffer (TBS) + 0.2% Tween® 20 at specified concentrations: Cell Signaling Phospho-Stat3 (Tyr705) Rabbit mAb (1:2000), b-actin (13E5) Rabbit mAb (1:2000), eBioscience IkB zeta (LK2NAP) Rat mAb (1:2000). Membranes were blocked with Odyssey blocking buffer TBS for 1hr and then incubated with primary antibody solution overnight at 4°C with gentle shaking. Membranes were washed with 1X TBS-T and subsequently incubated with the following secondary antibodies: 800CW Donkey anti-Rabbit IgG (1:10,000) or 800CW Goat anti-Rat IgG (1:10,000) in Odyssey Blocking Buffer (TBS) + 0.2% Tween® 20. Membranes were protected from light during incubation with secondary mixture for one hour at room temperature with gentle shaking. Membranes were subsequently washed with 1X TBS-T and imaged with the Odyssey® imaging system. Images were analyzed using ImageJ.

### qPCR

Reverse transcription of mRNA and subsequent qPCR was performed on RNA isolates of J774M cells treated with ENT (overnight) or STAT3i (72 hours) to determine changes in expression of genes associated with MDSC immunosuppressive function in J774M cells. Treated J774M cells were harvested and RNA was subsequently extracted using the RNeasy Mini Kit via manufacturer’s instructions (Qiagen, #74104). 1 μg of total RNA was used for reverse transcription via the SuperScript VILO cDNA Synthesis Kit (Invitrogen, #11754050) per the manufacturer’s instructions. qPCR of subsequent cDNA was performed using gene specific proprietary primer and probe constructs for selected genes obtained from TaqMan (ThermoFisher): *Stat3* (Mm01219775_m1); *Il10* (Mm01288386_m1); *Il6* (Mm00446190_m1); *Nox2* (Mm01287743_m1); *18s* (Mm04277571_s1). TaqMan Fast Advanced Master Mix (Applied Biosystems, #4444556) was used following manufacturer’s instructions. qPCR was performed on the StepOnePlus real-time PCR system (Applied Biosystems). The 2-step real-time PCR cycling conditions used were 50 °C for 2 min, 95 °C for 2 min, 40 cycles of 95 °C for 1 s, and then 60 °C for 20 s. Gene expression was quantified from raw data by calculating mean CT values from the three technical replicates and comparing the ΔCT values of each gene to the 18s reference gene. Fold changes were quantified as 2^ΔΔCT^.

### ChIP sample preparation and validation

ChIP-DNA was prepared using SimpleChIP Enzymatic Chromatin IP Kit #9005 (Cell Signaling Technology) following manufacturer’s protocol with sample-specific adjustments in micrococcal nuclease and sonication steps. 10X phosphate buffered saline (PBS) pH 7.4 from Quality Biological Inc. was used wherever PBS is indicated. Samples were digested by both micrococcal nuclease and sonication. This process was additionally optimized and followed by gel electrophoresis to ensure uniform shearing of DNA across the genome. Equal amounts of chromatin were used per IP step with exceptional performance (XP®) monoclonal antibodies validated for ChIP application (Cell Signaling Technology). Rabbit monoclonal antibodies were added in particular dilution based on an optimized concentration evaluated across a wide variety of commercial monoclonal antibodies. We used 1:50 diluted total H3 (4620) antibody as a positive control and 1:250 diluted Normal Rabbit IgG (2729) as a negative control. A 3527-5 Incubator Shaker (Lab-Line) was used during elution. ChIP-DNA was purified and measured following the ChIP kit protocol. The 1/50 portion (2%) of the same chromatin for each sample was used for DNA extraction skipping the antibody enrichment steps and was further used for qRT-PCR and sequencing as an input control. ChIP-DNA underwent qRT-PCR using a TaqMan® 7900HT Fast Real-Time PCR System (Applied Biosystems) per manufacturer’s recommendations. We used Johns Hopkins lab standard 10X PCR Buffer (20), dNTPs (Bioline), FAM (Thermo Fisher Scientific). Primers and probes designed in the promoter region of actively expressed GAPDH and RPL10 genes, and 3’ end of the transcriptionally repressed ZNF333 gene. Each sample was analyzed in triplicate and underwent one cycle of 10min at 95°C, and 50 cycles of 15s 95°C/60s 60°C. Relative fold enrichment of different histones in individual samples was quantified in triplicate relative to the 2% input sample using the 2-ΔΔCT method.

### ChIP Sequencing

Illumina’s TruSeq Nano DNA kit was used to prepare 12 samples. Sequencing was performed with Illumina paired end 150bp. Illumina’s CASAVA 1.8.4 was used to convert BCL files to FASTQ files using default parameters. Bowtie2 (v 2.2.5) was used for running the alignments to the mm10 genome using default parameters. MACS-2.1.0.2 was used for peakfinding using -g hs parameters. Overlaps were called using “intersectBed’’ utility of BEDTools. Motif-discovery was done using “MEME-chip”. deepTools was used to create a profile plot for scores over the transcriptional start site of all the genes.

### scRNAseq, quality control and analysis

For library preparation 10× Genomics Chromium Single Cell 3’ RNA-seq kits v2 were used. Gene expression libraries were prepared according to the manufacturer’s protocol. 20 processed tumor samples were sequenced in 4 batches Each batch had an equal assortment of samples from each treatment group to reduce technical biases. Illumina HiSeqX Ten or NovaSeq were used to generate ~6.5 billion total reads. Paired-end reads were processed using CellRanger v3.0.2 and mapped to the mm10 transcriptome with default settings. ScanPy v1.4 was used for quality control and basic filtering. For gene filtering, all genes expressed in less than 3 cells were removed. Cells expressing less than 200 genes or more than 8000 genes or have more than 15% mitochondrial gene expression were also removed. We implemented the Monocle3 v. 0.2.2.0 pipeline, including Batchelor for batch correction of library runs (Supp. Figure 5A, Haghverdi et al. 2018), UMAP low dimensional embedding, and Leidenbase 0.1.0 for community detection (implemented in Monocle3, Traag et al. 2019). Doublets were removed from final UMAP. For each cell type, Entinostat treated cells were compared to Vehicle cells, and each independent combination treatment was compared to Entinostat. To do this, the CDS subsets for each cell type were created and each treatment group comparison. fit_models function in monocle3 was then applied to those subsets for logistic regression analysis with library batches and treatment groups used as covariates in model (*model_formula_str = “~treatment + run”*). The beta estimate outputs were used to rank genes for pathway analysis and the p value statistics were FDR adjusted. Volcano plots were created using the enhanced volcano plot package v.1.7. SAVER v. 1.1.2 was used to impute missing data due to drop out, and ComBat function from the SVA package v.3.2 was used to batch correct the imputed scRNAseq expression matrix before generating heatmaps using ComplexHeatmap v2.42. The geneSetTest and barcodeplot functions from limma package v.3.44.1 were implemented for pathway analysis. Further psuedotime analyses were also performed with Monocle3 based on the Batchelor batch corrected UMAP embedding, and statistics from pseudotime were computed similarly to the treatment effects described above. InferCNV (of the Trinity CTAT Project. https://github.com/broadinstitute/inferCNV) package v. 1.4.0 generated a heatmap that illustrates inferred copy number variation across the cancer population in reference to fibroblasts, using pooled cells, the mm10 genome reference for chromosome mapping and default parameters. This confirmed that the NT2.5 cell line that was used to inoculate the mice, did not diverge across different tumors and treatments (Supp. Figure 5B). Of note, the consistent CNV observed in the tumor cells served as an internal control in that we would not have expected any significant shifts in tumor cells between mice and that any TME changes can be attributed to treatment alone and not any major discrepancies in the identity of the tumor cells. A small cluster of cells we called Cancer 2 which was consistent across treatments had a slightly different CNV profile and formed its own partition which seemed to be either a technical artifact or an uncharacterizable epithelial cell type and thus we removed from final UMAP (Supp. Figure 5B). Full code used in the analysis is made available on the following github repository: Dimitri-Sid/entinostat_ICI_scRNAseq

### Statistical analyses

For bar graphs, each dot represents a biological replicate, and each bar represents mean ± SEM. (n = 2-3 replicates/group). For Western blot, arginase activity, and qPCR data, significance was determined by a one-way ANOVA with Tukey’s multiple comparisons test. All experiments were repeated at least 3 times. For immunosuppression data, significance was determined by a two-way ANOVA with Dunnett’s multiple comparison’s test. All statistical analysis listed here were performed using GraphPad Prism v7.00. Statistically significant p values are abbreviated as follows: *p<0.05, **p<0.01, ***p<0.001, ****p<0.0001.

## LIST OF ABBREVIATIONS

scRNAseq: Single Cell RNA Sequencing
BC: Breast Cancer
TNBC: Triple Negative Breast Cancer
TME: Tumor Microenvironment
HDACi: histone deacetylase inhibitor
M-MDSCs: Monocytic Myeloid Derived Suppressor Cells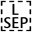
G-MDSCs: Granulocytic Myeloid Derived Suppressor Cells
DCs: Dendritic Cells
TAMs: Tumor Associated Macrophages
V: Vehicle (control, no treatment)
RTK: Receptor Tyrosine Kinase
E / ENT: Entinostat treatment
EP: Entinostat + anti PD1 treatment
EC: Entinostat + anti CTLA4 treatment
EPC: Entinostat + anti PD1 + anti CTLA4 treatment
CNV: Copy Number Variation

## DECLARATIONS

### Availability of data and materials

All scRNAseq raw and processed data files are available under NCBI BioProject accession number *PRJNA683665.*

### Ethics approval and consent to participate

All animal studies were approved by the Institutional Review Board of Johns Hopkins University.

## Acknowledgements

We would like to thank all members of the Jaffee lab for help throughout the course of these experiments. Additionally, we would like to thank Syndax Pharmaceuticals for supplying the entinostat, and AstraZeneca and Ionis for supplying STAT3i used in these experiments. We would like to thank the members of the Sidney Kimmel Comprehensive Cancer Center Experimental and Computational Genomics Core (ECGC), supported by NIH/NCI grant P30CA006973, for support with next generation sequencing and data processing.

## Author contributions

DNS, CR, BC, ERT, EJF, EMJ, LK prepared manuscript. ERT, BC, CR conducted murine, *ex vivo* and *in vitro* experimentation. DNS, LG, EJF conducted computational analyses. All authors reviewed manuscript.

---

**Supplemental Figure 1.**
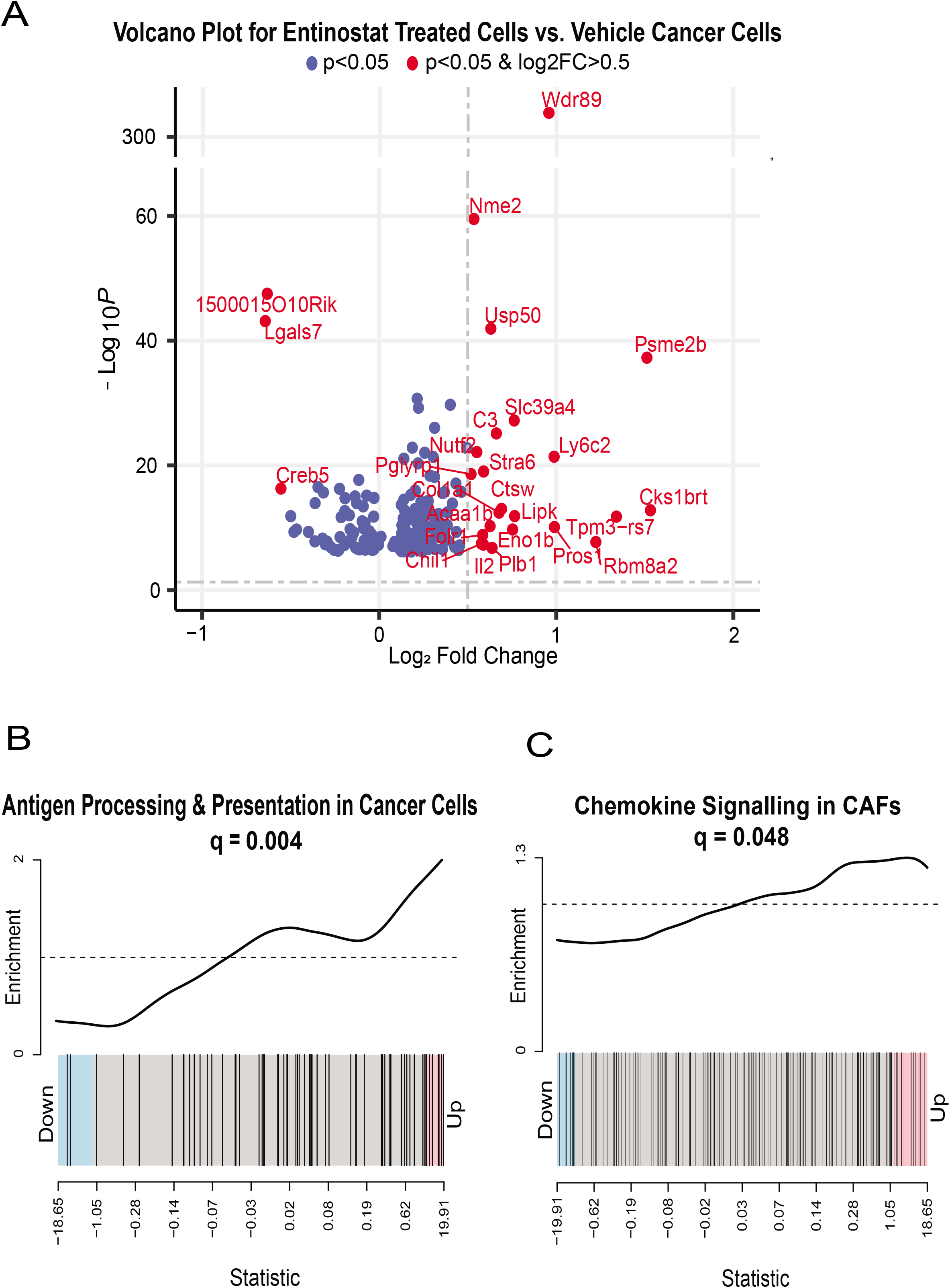
Differential expression analysis of non-immune populations. **A**, Volcano plot showing in red genes upregulated or downregulated with entinostat by at least 0.5 log2 Fold Change. **B**, Barcode plot showing that KEGG Antigen Processing and Presentation is significantly upregulated with entinostat in cancer cells. **C**, Barcode plot showing that KEGG Chemokine signaling pathways are significantly upregulated in CAFs with entinostat.

**Supplemental Figure 2.**
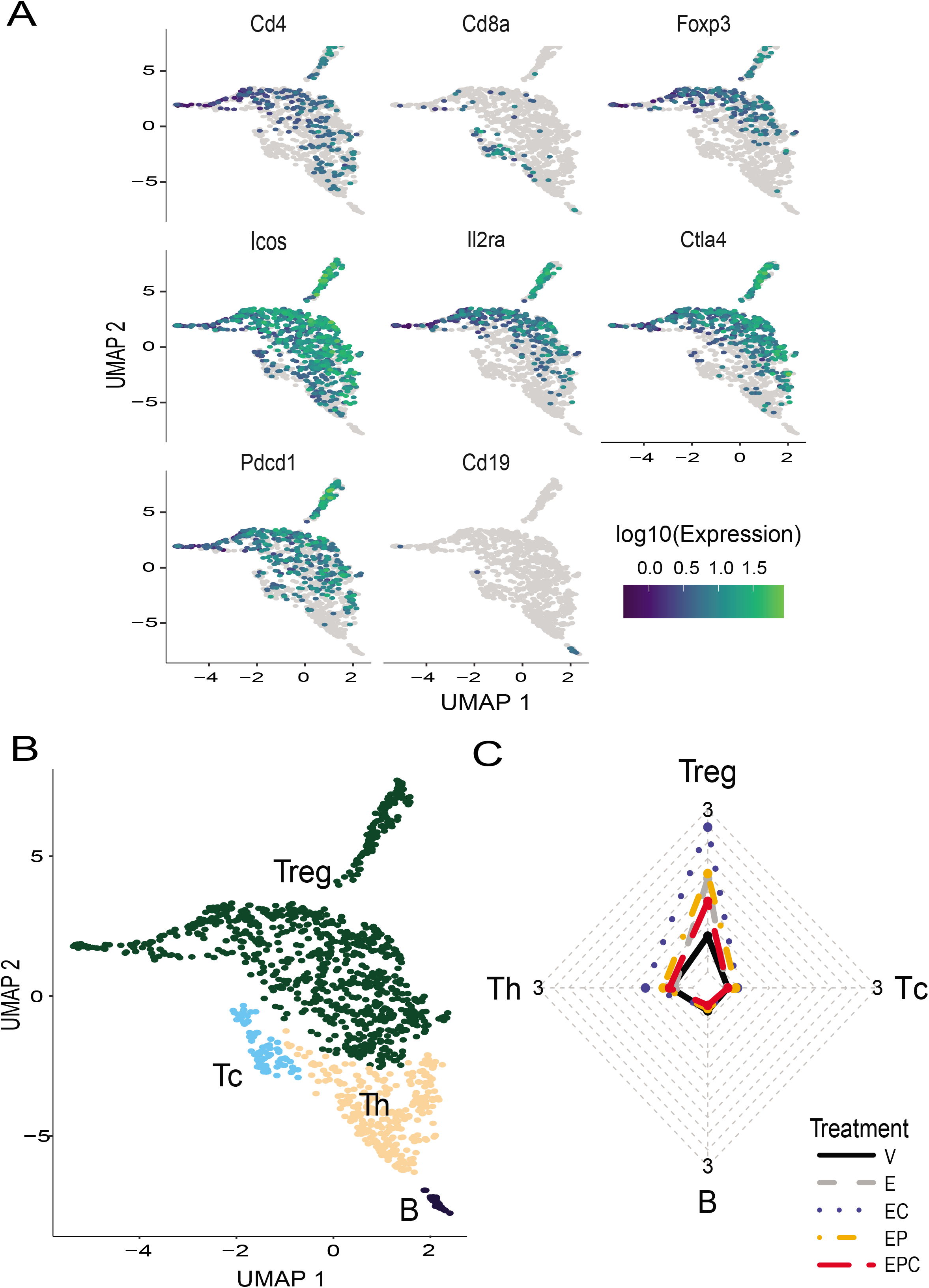
Tregs *(Cd4^+^ Foxp3^+^ Ctla4^+^* and T helpers *(Cd4^+^, Foxp3^-^)* are more abundant in combo Treatments. **A,** UMAPs of representative lymphoid cell subtype markers. **B**, Annotated clusters projected onto the UMAP of the lymphoid subset population. **C**, Radar plot of lymphoid cell type composition normalized to the total cell count from each group.

**Supplemental Figure 3.**
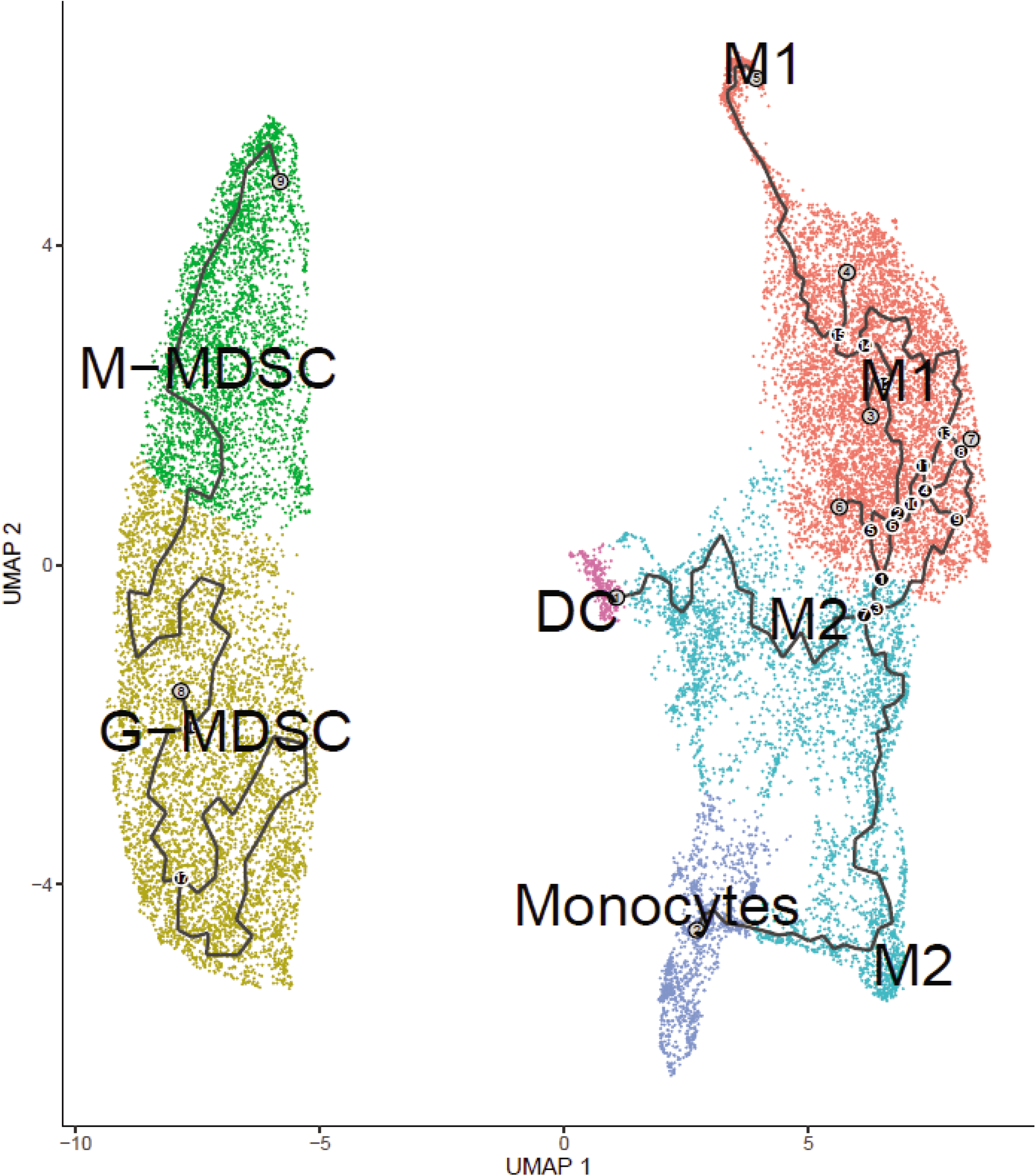
Pseudotime trajectory analysis of the myeloid cell populations. Trajectories are indicated by black paths.

**Supplemental Figure 4.**
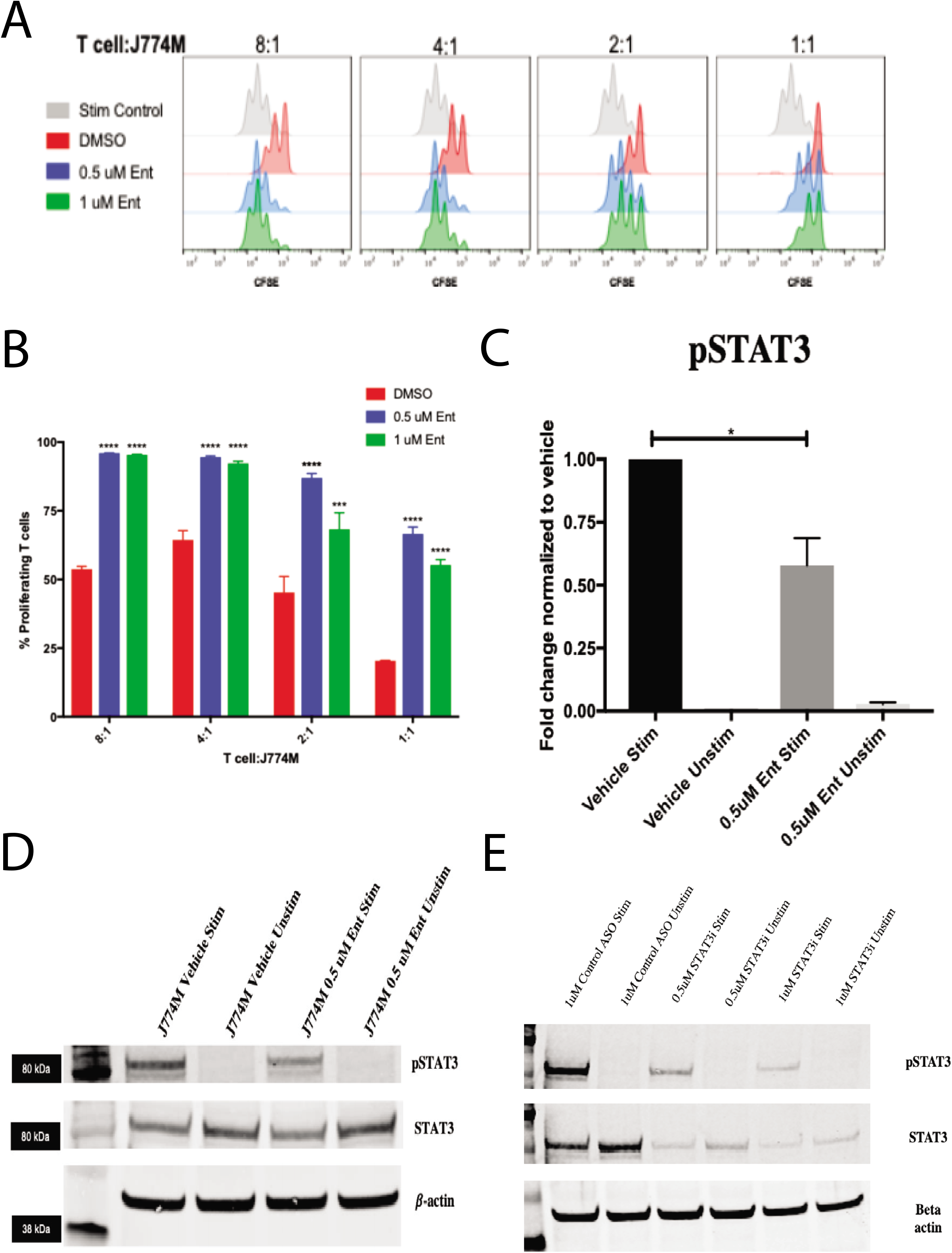
**A,** CFSE peaks of live CD8+ T cells co-cultured with ENT-pre-treated J774M cells. T cells were isolated from spleens of Balb/c mice and CFSE labeled, stimulated with Anti-CD3/CD28 beads and allowed to proliferate for 52 hours before harvest and analysis via flow cytometry. Each peak represents a cell division as determined by CFSE dilution. **B**, Quantification of percentages of proliferating CD8+ T cells with each treatment group compared to the DMSO control. n = 2 replicates per group. Each bar represents the mean±SEM. **C** and **D**, Western blot and corresponding quantification of J774M cells lysed and probed for phospho-STAT3, total STAT3, and beta-actin (n=2 biological replicates/group). **E**, Western blot of J774M cell lysates after treatment with STAT3i ASO probed for phospho-STAT3, total STAT3, and beta-actin (n=3 biological replicates/group).

**Supplemental Figure 5.**
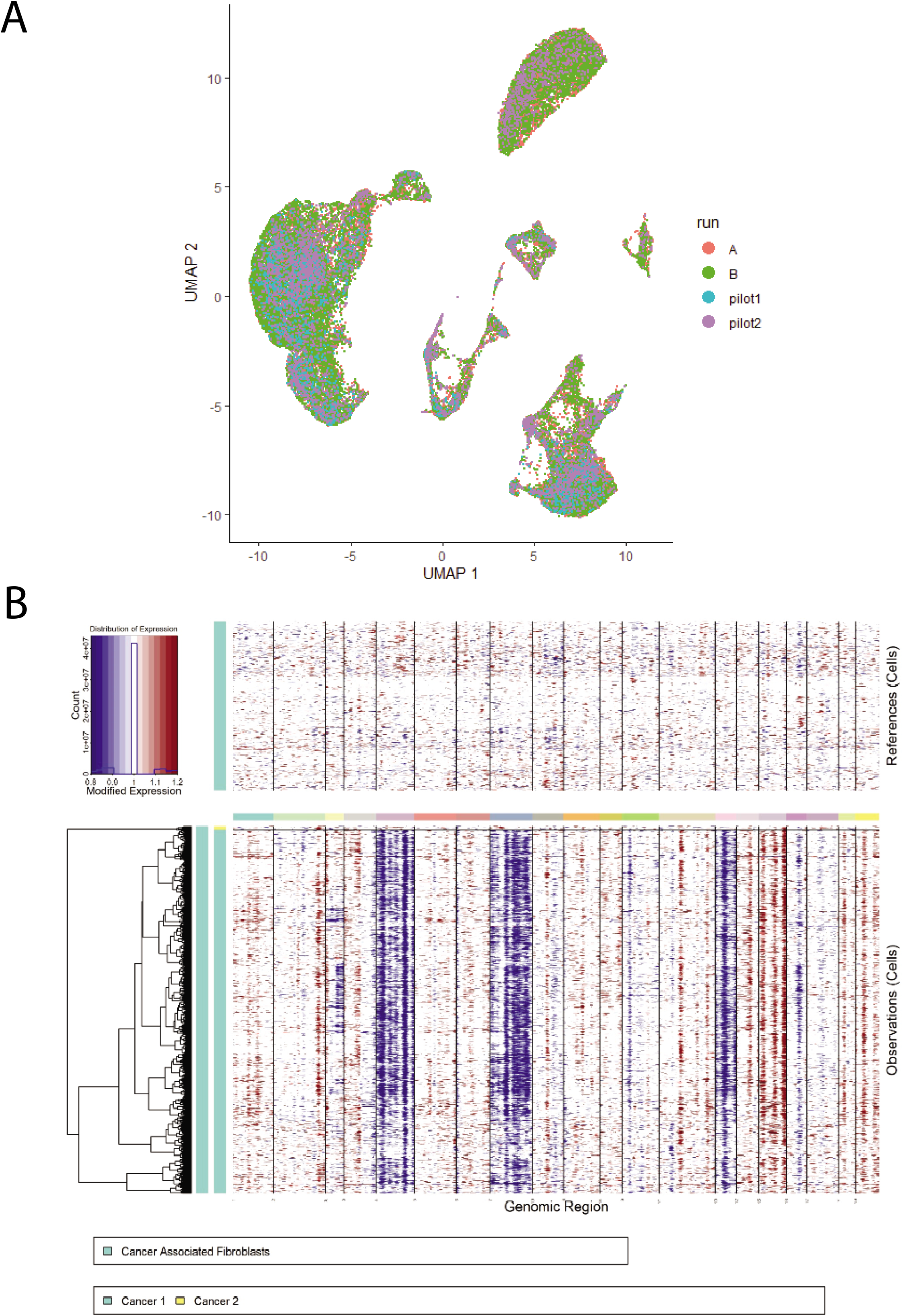
**A,** Batch-corrected embedding after applying Batchelor on sequencing runs. **B,** InferCNV output showing predicted CNV of the cancer cells when using CAFs as reference.

**Supplemental Table 1.**
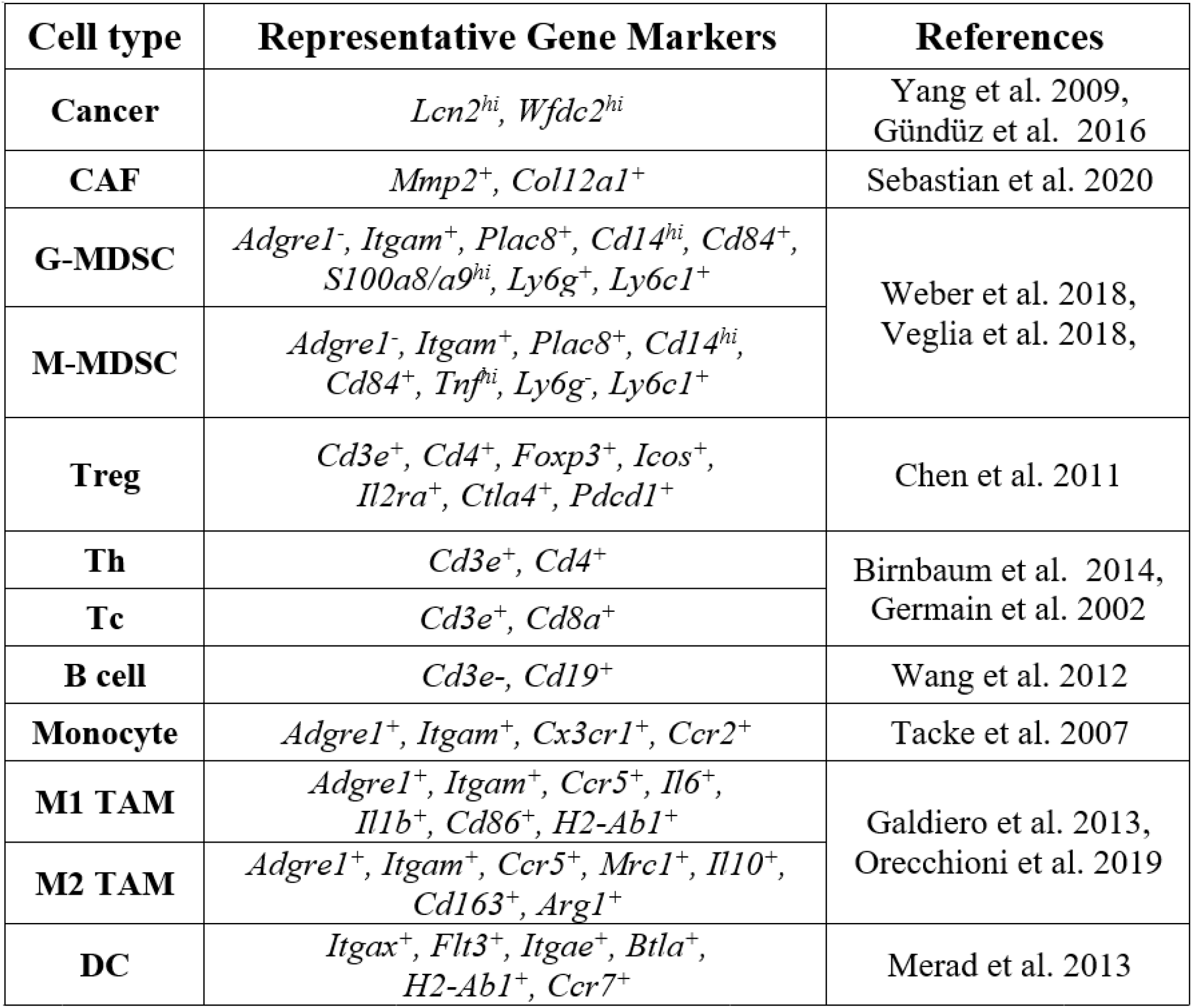
Representative gene markers used in this study.

**Supplemental Table 2.**
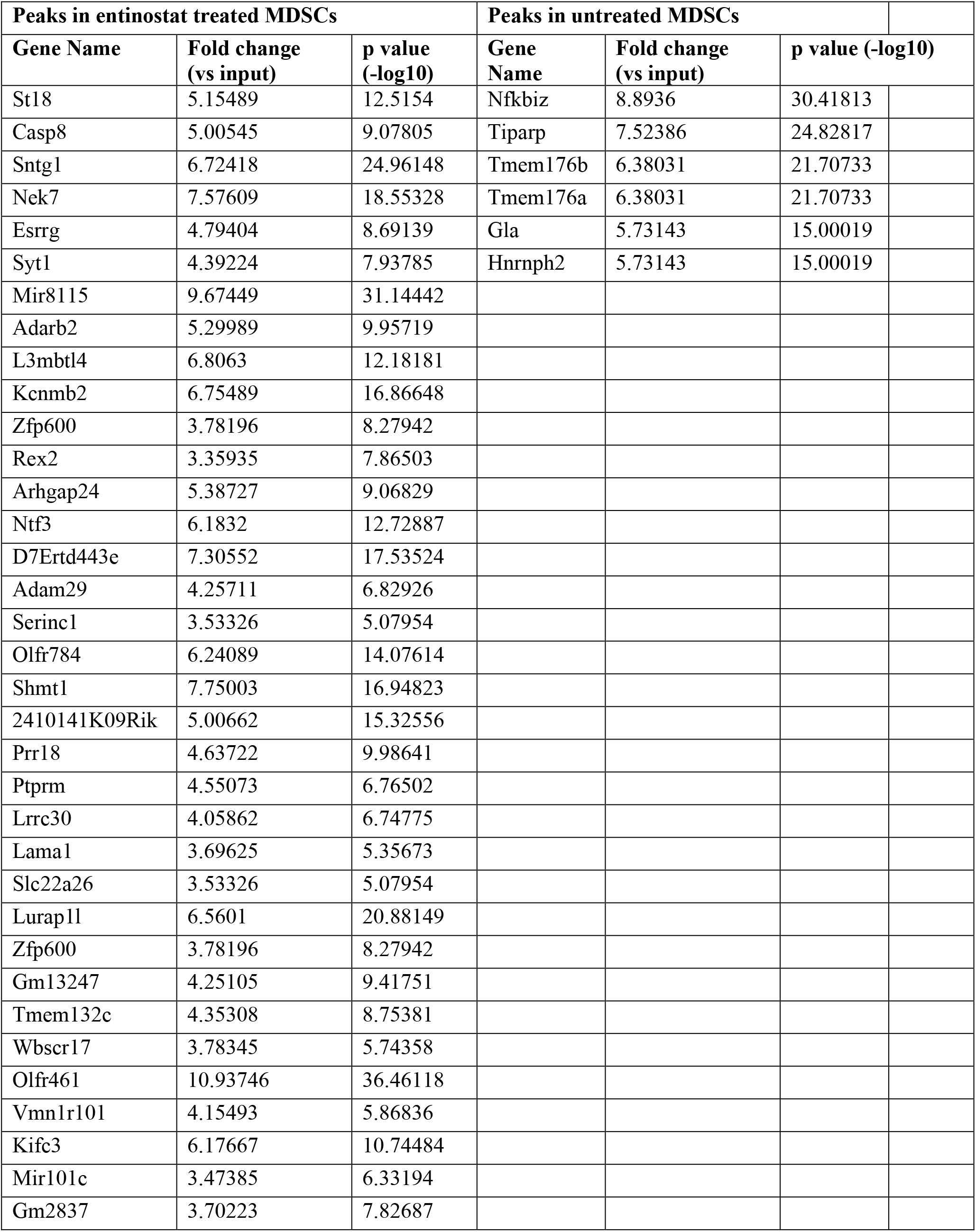
Summarized ChIP-seq results from entinostat-treated and untreated MDSCs.

